# Attractor-state itinerancy in neural circuits with synaptic depression

**DOI:** 10.1101/715532

**Authors:** Bolun Chen, Paul Miller

## Abstract

Neural populations with strong excitatory recurrent connections can support bistable states in their mean firing rates. Multiple fixed points in a network of such bistable units can be used to model memory retrieval and pattern separation. The stability of fixed points may change on a slower timescale than that of the dynamics due to short-term synaptic depression, leading to transitions between quasi-stable point attractor states in a sequence that depends on the history of stimuli. To better understand these behaviors, we study a minimal model, which characterizes multiple fixed points and transitions between them in response to stimuli with diverse time- and amplitude-dependences. The interplay between the fast dynamics of firing rate and synaptic responses and the slower timescale of synaptic depression makes the neural activity sensitive to the amplitude and duration of square-pulse stimuli in a non-trivial, history-dependent manner. Weak cross-couplings further deform the basins of attraction for different fixed points into intricate shapes. Our analysis provides a natural explanation for the system’s rich responses to stimuli of different durations and amplitudes while demonstrating the encoding capability of bistable neural populations for dynamical features of incoming stimuli.

## 1 Introduction

Mounting evidence suggests that neural ensembles can give rise to states of activity that are stable and attractor-like over a short period[1, 2, 3, 4, 5, 6, 7, 8]. However, given the range of timescales of neural processes, either slower processes or intrinsic noise typically ensure that an activity state does not remain stable for more than a few hundred milliseconds, even when a stimulus is constant. For example, when viewing images that can give rise to bistable percepts, a switching between the distinct perceived images arises [2, 4, 6, 7]. A similar switching can arise with auditory stimuli [1]. Analysis via Hidden Markov modeling [9, 10, 11, 12, 13, 14, 15] or change-point methods [16, 17] has suggested such state-switching in neural activity in sensory and decision-related tasks. Modeling work has shown how discrete attractor states can arise, and how either noise [18, 19, 20, 21, 22], or slow adaptation-like processes such as synaptic depression [23, 24], or a combination of the two [25, 26], can lead to transitions between these states, which we refer to as quasi-stable attractor states [8].

In this article we focus on how short-term synaptic depression [27, 28, 29] can lead to the instability of one quasi-stable attractor state, inducing a transition to a new state, which itself may be stable or quasi-stable. Mathematically, if one fixes the amount of synaptic depression by setting a slow, synaptic depression variable, to a constant, groups of neurons with strong self-feedback can possess multiple stable discrete attractor states. The system can resemble a relaxation oscillator with sufficiently strong depression and feedback [30], as the depression variable slowly decreases for an active group of neurons, reducing the within-group effective excitatory coupling until the activity can no longer be maintained. Once inactive, the depression variable slowly recovers, allowing for connections to re-strengthen and activity to recommence. In other ranges of parameters, the remnant of such potential oscillatory behavior leads to a rich repertoire of states and state transitions in response to simple stimuli when the stable states of such systems are fixed points.

We characterize such systems with small numbers of (one to five) potentially bistable groups of neurons, via the number of fixed points and their basins of attraction. We assess how different fixed points are reached as a function of the amplitude or duration of stimuli, as well as the system’s state before stimulus onset. In particular, we use an extended Wilson-Cowan model [31]and incorporate synaptic depression, to show how weak coupling between distinct bistable populations impacts the states’ basins of attraction, which can be deformed into complex shapes. In so doing, we offer an initial explanation of the rich information processing capabilities of high-dimensional networks with multiple attractor states and slow synaptic dynamics [32].

The rest of this paper is organized as following: We introduce the rate model with synaptic depression in Sec. 2, and derive the dimensionless form that will be used for later analysis. Section 3 focuses on a single population’s dynamics. We derive the condition for bistability and analyze the linear stability of fixed points for the single population. We find that the unit’s responses to stimuli exhibit history dependence and provide an explanation using a reduced 2D model, which is derived by assuming a separation of time-scales. In Sec. 4, we consider two coupled populations and obtain the basins of attraction for attractors as well as bifurcations with cross-couplings. Similarly to the behavior of a activity in a single population model, dynamical responses to stimuli can traverse multiple basins of attraction, leading to stimulus-duration dependence as well as history dependence and stimulus-amplitude dependence of the final state. Section 5 extends the discussion to multiple populations with weak and random cross connections. We numerically explore how the number of states scales with the connectivity and depends on initial conditions. We summarize these results in Sec. 6 and discuss some open questions for future research.

## 2 Model

We consider a network of *N* neural populations, each of which can be characterized by its mean firing rate, *r_i_*. The dynamics of the population rate, *r_i_*, in response to time-varying current, *I_i_*(*t*), is given by a generic form,

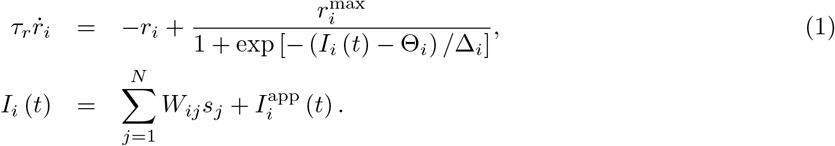

Here 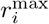 is the maximum firing rate, Θ_*i*_ is the input threshold for the half-maximum firing rate, and Δ_*i*_ is inversely proportional to the slope of the input-output curve. The input current, *I_i_*(*t*), consists of two parts: 1) synaptic currents from the network with a connectivity *W_ij_*, which quantifies the coupling strength from population *j* to population *i*; 2) an applied current 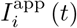.

The time-varying effective synaptic input, *s_i_*, arising from a population *i*, is given as a fraction of the maximum possible (so *s_i_* ∈ [0,1]). We assume spikes are emitted from the population via a Poisson process and include a short-term synaptic depression factor, *d_i_* ∈ [0,1], with 0 indicating a fully depressed synapse. With these assumptions, the mean dynamics of *s_i_* and *d_i_* take the following form [24],

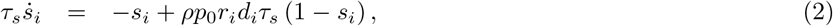

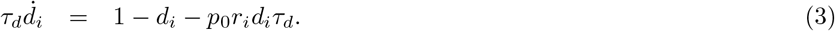

The parameter *p*_0_ gives the fraction of docked vesicles released per spike. *ρ* is the fraction of open receptors bound by maximal vesicle release, such that *ρp*_0_*d_i_* is the fraction of closed synaptic receptors that open, so is proportional to the increase in the synaptic current for a given presynaptic spike.

The time constants for the mean firing rate, the synaptic current, and the depression variable are denoted respectively as *τ_r_*, *τ_s_*, and *τ_d_*. Since these dynamical variables vary over distinct time scales, it is convenient to rescale the time and to normalize the rate: *t*/*τ_r_* → *t*, 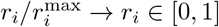, as well as to scale the input and threshold by 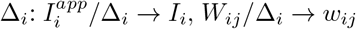, and Θ_*i*_/Δ_*i*_ → *θ_i_*. The dimensionless equations then become

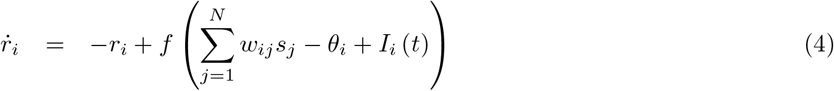

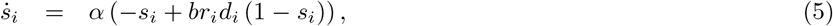

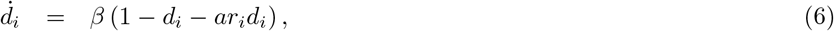

where *f*(*x*) = (1 + *e*^−*x*^)^−1^ is the logistic function, *θ_i_* is the activation threshold. The weight matrix *w_ij_* determines the coupling strengths within a unit and between units. Two remaining time-scales are characterized by *α* = *τ_r_*/*τ_s_* and *β* = *τ_r_*/*τ_d_*. In this paper, we assume that the short-term depression varies over a slow time scale compared with the firing rate and the synaptic current. This situation arises when the timescale for recovery from depression is significantly longer than other time constants, such that *τ_d_* ≫ *τ_s_* > *τ_r_*. For example, we set *τ_d_* = 250ms, *τ_s_* = 50ms, and *τ_r_* = 10ms in simulations, and *p*_0_ = 0.5. The dimensionless parameters

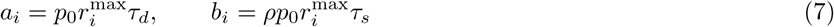

quantify the degree of synaptic depression and the amplitude of synaptic currents respectively. With slow depression, *a_i_* ≫ *b_i_*.

Finally, all cell-groups are assumed to be comprised of neurons with identical parameters. For most simulations we choose the standard parameter set: *a* = 6.25, *b* = 1.25, *w_ii_* = 40, and *θ_i_* = 5, unless noted otherwise. In a control scenario [see Fig. 4(c)], to demonstrate the importance of synaptic depression, we produce a network without depression by setting *τ*_d_ → 0 thus *a* → 0 and *d_i_* → 1. Then the firing rate is solely driven by the synaptic current within a time window of *τ_s_*.

## 3 Single Population

### 3.1 Multiple fixed points

For a single population, its dynamics are governed by

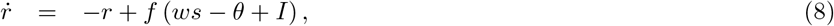

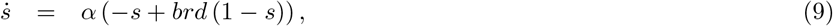

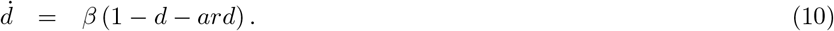

At a fixed point 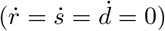, the solution to Eqs. (8)–(10) satisfies

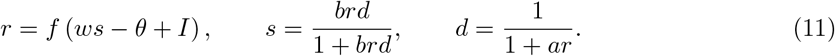

Since the logistic function *f* is invertible, we define *g*(*r*) = *f*^−1^(*r*) = ln [*r*/(1 ‒ *r*)]. Then at a fixed point, one must have

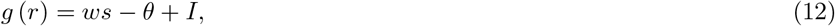

where *s* is parameterized by *r* as *s*(*r*) = *br*/(1 + (*a* + *b*)*r*). Solving for a fixed point is equivalent to finding an intersection between two curves *g*(*r*) = ln [*r*/(1 – *r*)] and *ws*(*r*) – *θ* + *I*. Since both *g* and *s* are monotonically increasing and *s* is positive-definitive (*s*′(*r*) > 0), there is at least one solution.

Figure 1 (a) and (b) show two cases with one and three fixed points. The tangency condition in Fig. 1(a) can be expressed as *g*′(*r*) = *ws*′(*r*), which is equivalent to a parametric equation for *w*:

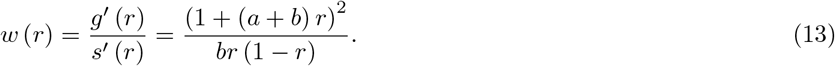

**Figure 1.**
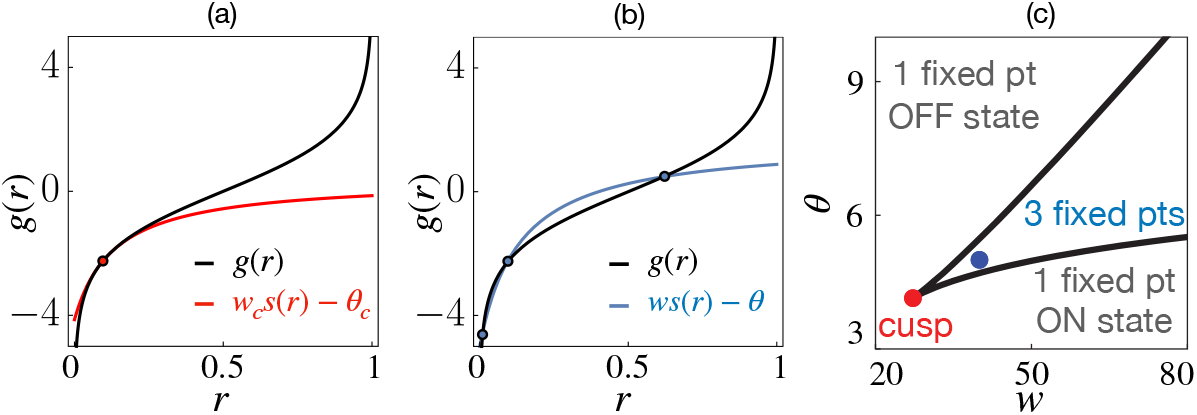
(a)-(b): A plot of *g*(*r*) = ln (*r*/(1 – *r*)) together with *ws*(*r*) – *θ* provides a graphic method for finding fixed points (filled circles, where the two lines cross) of a single neural population in the absence of external stimuli (*I* = 0). The parameter set for the self-coupling weight *w* and the input threshold *θ* are indicated in panel (c) with a red dot (for parameters of Fig. 1a) and a blue dot (for parameters of Fig. 1b). (c): The solid curve separates the region with three fixed points from the region with one fixed point in the *w-θ* plane. With *w* = 40 and *θ* = 5 (blue dot) the population is bistable. The cusp point (red dot) is located at (*w_c_*, *θ_c_*) = (27.2,4.14) according to Eqs. (15) and (16).

This equation describes the onset of a bifurcation and delineates a region of parameters with a single fixed point from a region with multiple fixed points. Likewise, by rewriting Eq. (12), we find a parametric equation for *θ*,

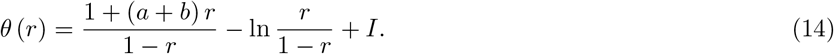

Graphing Eqs. (13)–(14) in the *w*-*θ* plane [Fig. 1(c)], systems with three fixed points arise from parameters within the resulting wedge-shaped region, while systems with one fixed point arise from parameters outside that region. Similar wedge boundaries were found in [33] for rate models without synaptic depression. At a cusp point (*w_c_*, *θ_c_*), *w*′(*r_c_*) = 0; A co-dimension-2 bifurcation takes place. Solving this condition using Eq. (13) yields *r_c_* = (*a* + *b* + 2)^−1^ and from Eqs. (13) - (14), the location of the cusp point is:

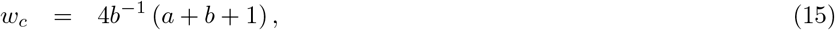

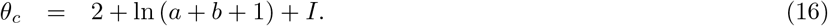

Dimensionless parameters *a* and *b* are chosen as *a* = 6.25 and *b* = 1.25. With *w* = 40 and *θ* = 5 (blue dot) the population is bistable. The cusp point (red dot) is located at (*w_c_*, *θ_c_*) = (27.2,4.14) according to Eqs. (15) and (16).

Equation (7) shows that with slow depression *a* ≫ *b*, so *w_c_* ≫ 1. Namely, the existence of multiple fixed points requires strong self-excitation. For a given *w*(*w* > *w_c_*), the width of this region (measured by the input threshold) is

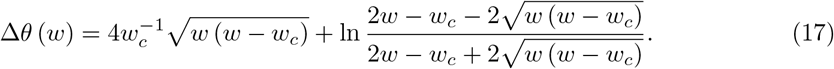

With strong self-coupling, *w* ≫ *w_c_*, the width scales linearly with *w*, 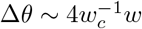, indicating that multiple fixed points are robust under a wide range of stimuli.

### 3.2 Linear stability and bifurcations

The stability of a fixed point (*r*, *s*, *d*) is captured by the Jacobian matrix via linearizing Eqs. (8)–(10),

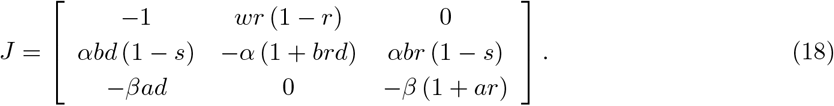

Here we have used the identity *f*′ = *f*(1 – *f*) = *r*(1 – *r*) at the fixed point. From Eq. (11), *s* and *d* can be further expressed in terms of *r*. Eigenvalues, λ, of the 3 × 3 Jacobian matrix, are roots of a cubic characteristic polynomial, *P*(λ) = λ^3^ + *A*_2_λ^2^ + *A*_1_λ + *A*_0_ with coefficients

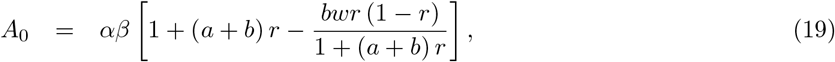

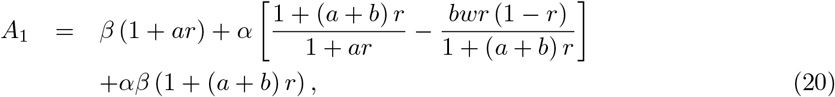

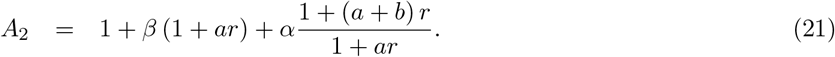

Using the Routh-Hurwitz criterion [34], the fixed point is stable (Reλ < 0) if *A*_0_, *A*_2_ > 0 and *A*_1_*A*_2_ – *A*_0_ ≡ *H*_2_ > 0. When an eigenvalue becomes zero (*A*_0_ = 0) or purely imaginary (*H*_2_ = 0 and *A*_0_, *A*_2_ > 0), the fixed point undergoes a saddle-node (SN) or a Hopf bifurcation (HB).

From Eq. (19), the condition for a saddle-node bifurcation (*A*_0_ = 0) implies [1 + (*a* + *b*)*r*]^2^ = *bwr* (1 – *r*), which is the same as Eq. (13). On the other hand, since *A*_2_ is always positive, the condition for a Hopf bifurcation (*A*_1_*A*_2_ = *A*_0_ > 0) converts into

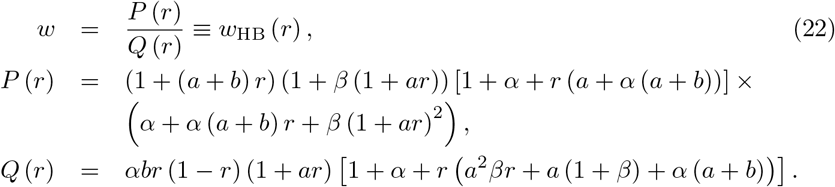

If we fix the threshold *θ* and view the external stimuli *I*_app_ as another bifurcation parameter, we obtain a parametric equation from Eq. (12),

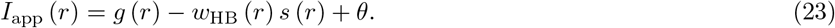

Figure 2 is the bifurcation diagrams of a single population. In Fig. 2(a), the steady-state (characterized by the synaptic current *s*) changes stability as applied stimuli *I*_app_ vary (*w* and *θ* are fixed). There are four bifurcations: For large inhibitory stimuli (*I*_app_ < *I*_SN_1__ = −0.4627), the low firing rate state (OFF state) is stable. When *I*_app_ = *I*_SN_1__, the high firing rate state (ON state) and a saddle point emerge from a saddle-node (SN) bifurcation. The ON state is unstable till a subcritical Hopf bifurcation (HB) takes place. Then both the ON and OFF states become stable with an unstable limit cycle surrounding the ON state. The size of the limit cycle (red open circles) grows and merges with the saddle point via a saddle-homoclinic orbit (SHO). The population remains bistable between the Hopf and another SN bifurcation (*I*_SN_2__ = 0.3002). When *I*_app_ > *I*_SN_2__, the ON state is the system’s only fixed point. The 2d model (blue open circles) is derived from slow-fast separation and will be discussed in Section 3.4.

**Figure 2.**
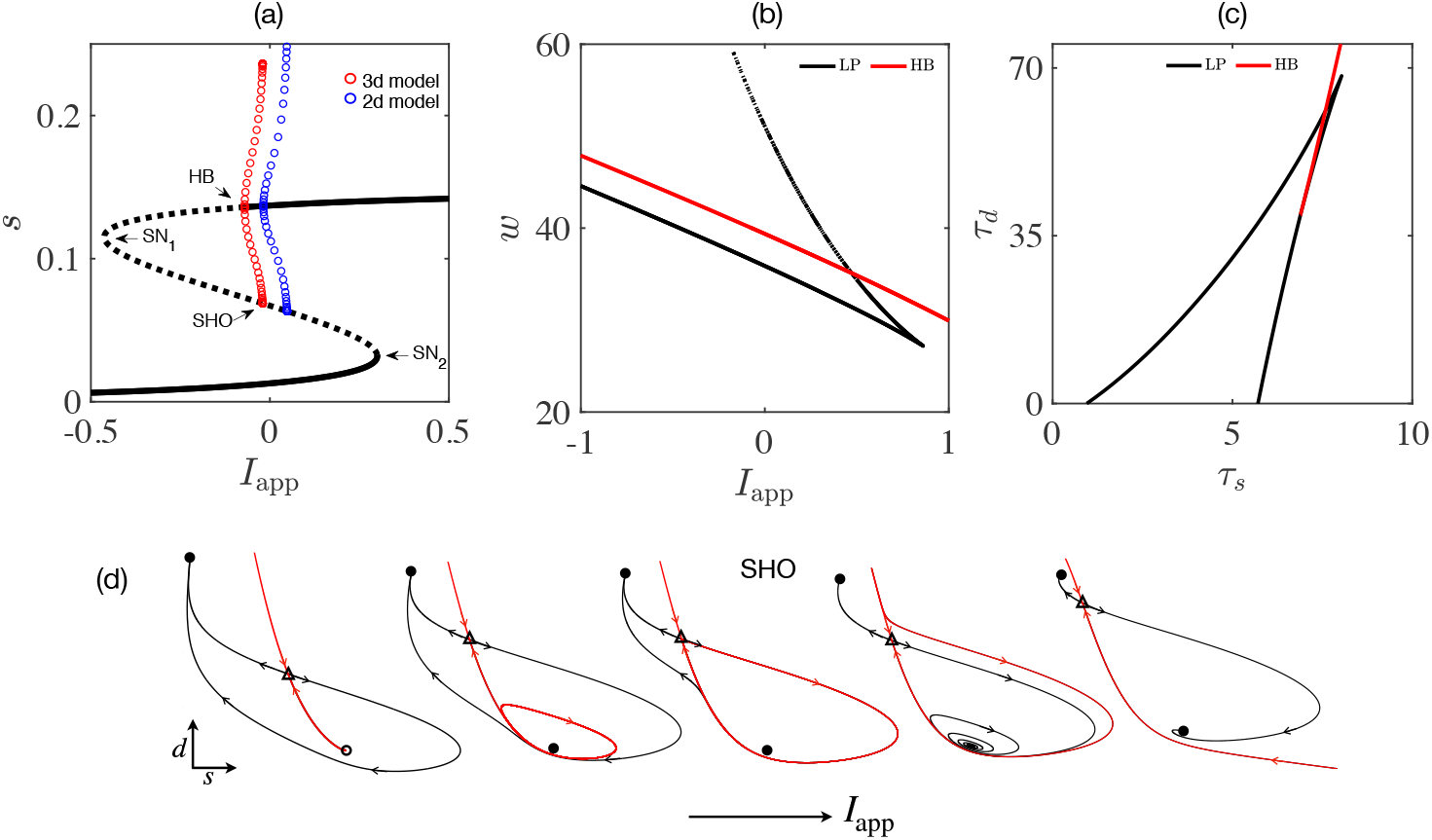
Bifurcation diagrams for a single population. (a) Bifurcations of the synaptic current, *s*, against stimulus current, *I*_app_. Two saddle-node (SN) bifurcations take place at *I*_SN_1__ = −0.4627 and *I*_SN_2__ = 0.3002. An unstable limit cycle arises from a subcritical Hopf bifurcation (HB) at *I*_HB_ = {−0.07069, −0.01817} for the 3d (red) and 2d (blue) model (see Section 3.4). The limit cycle merges with the saddle in a saddle-homoclinic orbit (SHO) at *I*_SHO_ = {−0.02013, 0.04802}. The system is bistable between HB and SN_2_. (b) Two-parameter bifurcations with stimuli *I*_app_ and self-coupling weight *w*. The bistable region is between the top limit point (LP) line and the HB line. (c) Two-parameter bifurcations with time constants *τ_s_* and *τ_d_* (both measured in *τ_r_*). The bistable region is between the left LP line and the HB line. A cusp point is at (*τ_s_*, *τ_d_*) ≈ (8.0, 68.1). (d) Bifurcation sequence in the *s-d* plane between SN_1_ and SN_2_. The OFF state remains stable (filled circle with small *s* and large *d*). Red (black) lines are stable (unstable) invariant manifolds of the saddle (triangle). The ON state is initially unstable (open circle) and gets stabilized at the Hopf bifurcation. The central plot depicts how the unstable limit cycle terminates at the saddle when *I*_app_ = *I*_SHO_.

In Fig. 2(b), using Eqs. (22) and (23), we plot the HB curve as well as the SN curves in the *I*_app_-*w* plane. The bistable region is above the HB curve and below the upper boundary of the SN curve. The wedge region ends at a cusp point (*w_c_*, *I_c_*) = (4*b*^−1^ (*a* + *b* + 1), *θ* – ln(*a* + *b* + 1) – 2).

Since parameters *a* and *b* depend on time constants *τ_s_* and *τ_d_*, it would be interesting to see how their change affects the fixed points’ stability. In Fig. 2(c), we numerically compute the bifurcation diagram with varying time constants *τ_s_* and *τ_d_*, which has similar structure to that in Figs. 1(c) and 2(b). The region between the left boundary of the SN curve and the HB curve supports bistable solutions. Hence the bistability is robust for slow depression and a wide range of parameters.

Figure 2(d) shows stable (red lines) and unstable manifolds (black lines) of the saddle point (a triangle) in the *s-d* plane between the two saddle-node bifurcations. When *I*_SN_1__ < *I*_app_ < *I*_HB_ (first plot from the left), the ON state is unstable (labeled as an open circle), the basin of attraction for the OFF state is the whole plane. At the Hopf bifurcation, the ON state becomes stable and has a small attracting basin bounded by the unstable limit cycle (the red loop). Right after the limit cycle merges with the saddle’s unstable manifold (*I*_app_ ≥ *I*_SHO_), the two stable manifolds (red lines) are initially very close to each other then split apart. One of them orbits around the ON state before heading back to the saddle point. The basin of attraction for the ON state grows and exceeds that of the the OFF at larger *I*_app_. The changing basins contain essential information for understanding responses to constant stimuli.

### 3.3 Responses to stimuli

In the previous section, we have seen that a single neural population is near several bifurcations. The neural activity thus exhibits rich dynamics in response to constant stimuli. Using the Heaviside function, an applied current with amplitude *I*_app_, duration *τ*_dur_ and onset time *t*_0_ can be written as

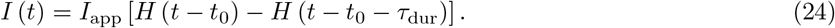

Following the end of the stimulus, the population’s firing rate converges to a stable fixed point. The final state, which may differ from the initial state, depends on the duration and amplitude of the stimulus, which together determine the system’s state at the time of stimulus offset. Figure 3 shows simulated responses to stimuli in three scenarios:

- Column (a): The population is initially in the OFF state, receiving stimuli with fixed amplitude (*I*_app_ = 0.45) and increasing duration (across rows 1 to 5). The population is OFF after a short stimulus (*τ*_dur_ = 20). For intermediate duration (*τ*_dur_ = 40), the population turns ON. With a longer stimulus (*τ*_dur_ = 60), the final state is OFF again. Even longer stimuli (*τ*_dur_ ≥ 80) keep the population in the ON state.
- Column (b): The population is initially in the OFF state, receiving stimuli with a fixed duration (*τ*_dur_ = 60) and increasing amplitude (across rows 1 to 5). The population is OFF after a weak stimulus (*I*_app_ = 0.3). For intermediate amplitude (*I*_app_ = 0.37), the population turns ON. With a stronger stimulus (*I*_app_ = 0.42), the final state is OFF again. Even stronger stimuli (*I*_app_ ≥ 0.6) keeps the population in the ON state.
- Column (c): The population is initially in the ON state, receiving stimuli first with a fixed amplitude (*I*_app_ = 0.8) and increasing duration (across rows 1 to 3), then with fixed duration (*τ*_dur_ = 10) and increasing amplitude (across rows 1 and 4 to 5). Similarly to the above two cases, the final state depends non-monotonically on the amplitude and duration of the stimulus.

**Figure 3.**
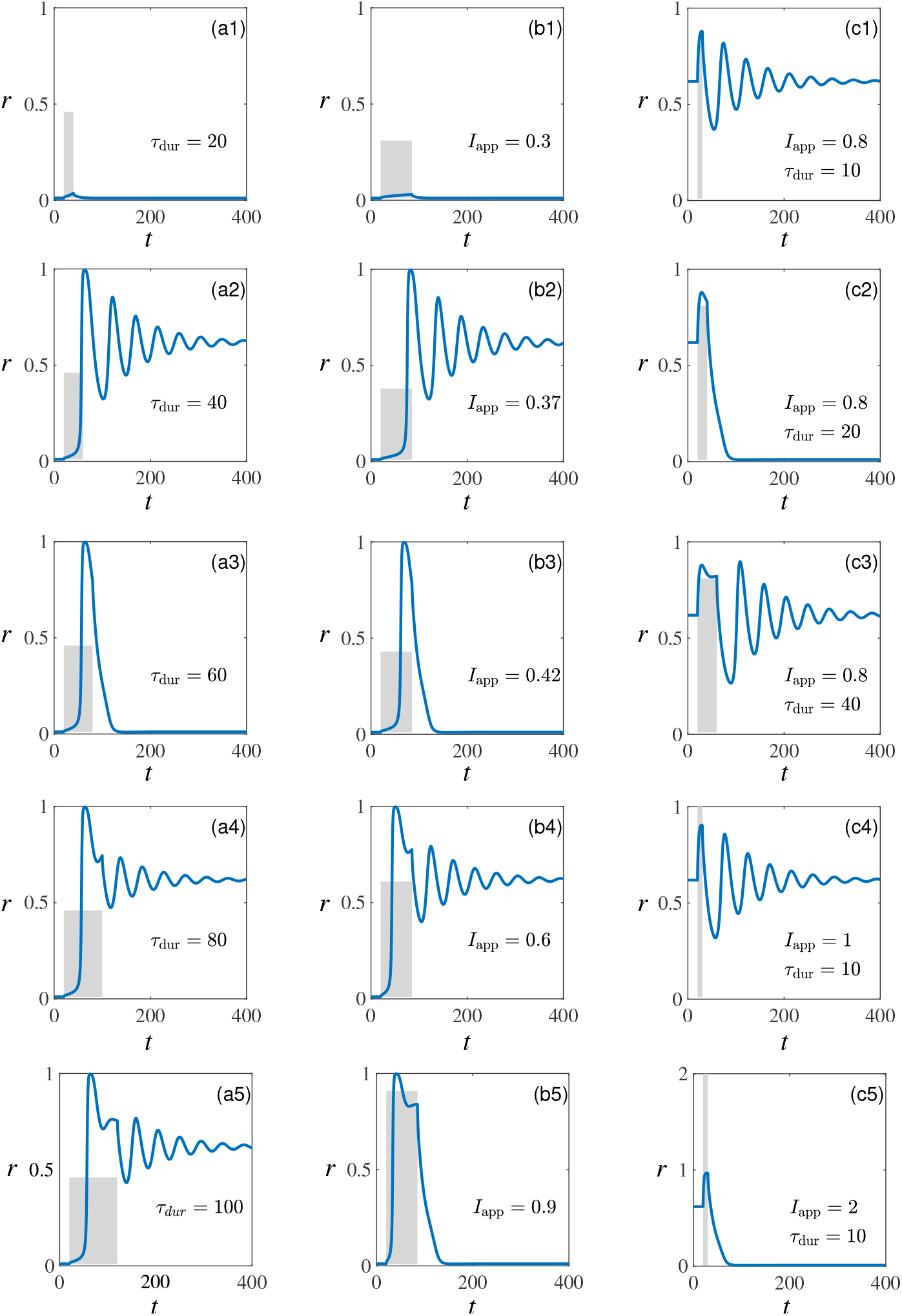
Responses of a single population to stimuli (shaded rectangles) show history-dependence. (a1) – (a5): State switches for the OFF state under stimuli with fixed amplitude (*I*_app_ = 0.45) and increasing durations; (b1) – (b5): State switches for the OFF state under stimuli with fixed duration (*τ*_dur_ = 65) and increasing amplitudes; (c1) – (c5): State switches for the ON state under stimuli with varying amplitudes, (c1) – (c3), and durations, (c1) and (c4) – (c5).

For other values of *τ*_dur_ and *I*_app_, the responses are summarized as phase diagrams in Fig. 4(a)-(b), which show history-dependence and reentrance behavior. Synaptic depression is essential for the non-monotonic (reentrant) behavior, as in its absence [Fig. 4(c)], the phase diagram for the final state has a rather simple boundary between the ON and the OFF states.

**Figure 4.**
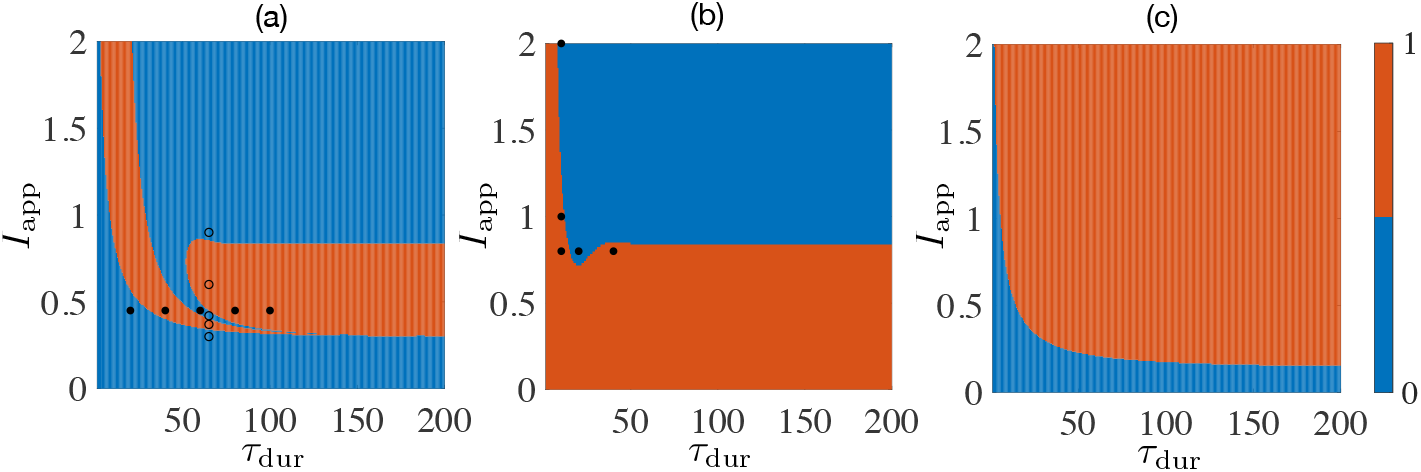
A single population’s final states as functions of stimuli’s duration *τ*_dur_ and amplitude *I*_app_. (a) The initial state is OFF. The filled and open circles indicate the stimuli used for columns a and b in Fig. 3. (b) The initial state is ON. Filled circles are values used in column c of Fig. 3. (c) Final states of a single population without synaptic depression (the initial state is OFF). The color coding is blue stands for the OFF state (labeled as 0) and red stands for the ON state (labeled as 1).

### 3.4 Separation of time scales

The phase diagram [Fig. 4(c)] in the last section indicates that synaptic depression is essential for the final state to depend non-monotonically on input. Since the depression takes place at a much slower time scale than that for changes in the synaptic current and the rate, *τ_d_* ≫ *τ_s_* > *τ_r_*, it is convenient to separate the fast dynamics from slower ones [35]. We can either fix the depression variable and solve for fixed points of the *r-s* subsystem [Eqs. (8),(9)], or we can assume the rate instantaneously reaches a steady state 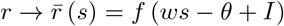 and get a reduced *s-d* subsystem [Eqs. (9),(10)].

In the first scenario [Fig. 5(a)], synaptic depression acts as a slow perturbation. When *d* = 1 (no depression), the population has two stable fixed points (the ON and OFF state) separated by a saddle. As the firing rate approaches the ON state, there is increasing synaptic depression (*d* decreases). At a critical value (*d_c_* ≈ 0.18), the ON state and the saddle annihilate in a saddle-node bifurcation and the population switches to the OFF state. In the second scenario, the firing rate is a function of the synaptic input. In the bistable region [Fig. 5(b)], the steady-state 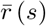 has a steep slope at small *s*. So the system is sensitive to stimuli and feedback. In Fig. 5(c)-(d), we compare solutions of the full system with the two reduced subsystems; In particular, evolutions of the synaptic current s and the effective firing rate *rd*.

**Figure 5.**
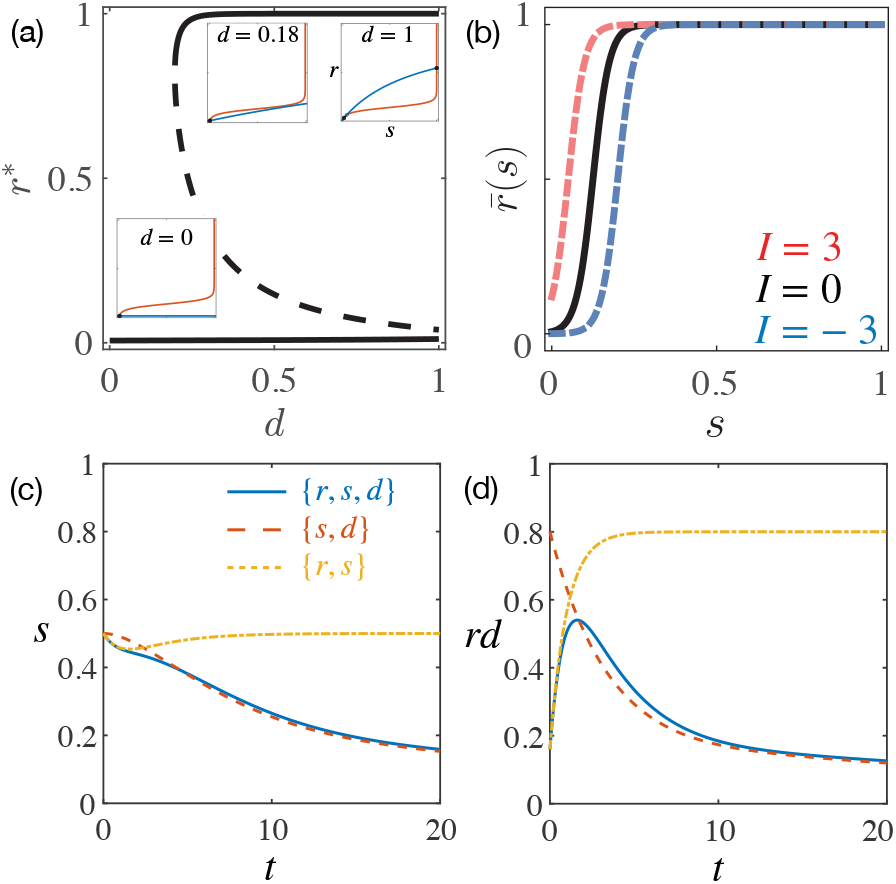
(a): Depression-induced bifurcations in the *r-s* subsystem. Solid (dashed) lines are stable (unstable) fixed points. Insets show fixed points as intersections between *r*-nullcline (red) and *s*-nullcline (blue). (b): The steady firing rate 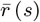 as a function of the synaptic input under three different stimuli. (c)-(d): Comparison between the full 3d system (solid blue line), the reduced *s-d* model (dashed red line), and the reduced *r-s* model (dotted yellow line) in terms of the synaptic current *s* and the effective firing rate *rd*.

In Fig. 5(c)-(d), the *r-s* subsystem captures the short-term behavior, whereas the *s-d* subsystem better approximates long-term evolution. Bifurcation analysis [Fig. 2(a)] reveals the same bifurcations in the *s-d* model as in the 3d system, other than minor numerical differences in critical values for the Hopf and SHO points. Therefore, phase portraits of the *s-d* model [like Fig. 2(d)] are useful for understanding the population’s responses to stimuli.

We note that in the reduced 2d model, there are still two time-scales (*s* vs. *d*). So the nullclines (not shown in the figure) and dynamics resemble a generic fast-slow system [36, 37]. Let us now return to Fig. 2(d) and walk through the bifurcation sequence.

With our standard parameter set, the system has a stable OFF state, a saddle point, and an unstable ON state for not-too-strong inhibitory stimuli (−0.5 < *I*_app_ < 0). A Hopf bifurcation takes place at *I*_HB_ = −0.07069, which stabilizes the ON state. The unstable limit cycle (born from the Hopf bifurcation) separates basins of attraction for the ON state and the OFF state. For small excitatory stimuli, the ON state’s basin is small. When *I*_app_ = *I*_SHO_ = −0.02013, the limit cycle crashes onto the saddle. Thus the system is already bistable at zero stimulus and remains bistable for small excitations. As the input increases, the ON state’s basin size enlarges rapidly. In the last graph in Fig. 2(d), the system is still bistable but the ON state’s basin extends to the entire region that is above the saddle’s stable manifolds (red lines).

For strong stimuli (*I*_app_ > 0.5), the OFF state annihilates with the saddle (not shown in the figure). The system is in the ON state. When the input current shuts off, the vector field (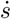 and 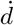) and basins of attraction all resume to the case with *I*_app_ = 0. Then the system’s instantaneous location in the *s-d* plane may be in the small basin of the ON state or the large basin of the OFF state. So the final state depends sensitively on the amplitude and the duration of a stimulus. In short, deformed basins of attraction under stimuli cause the history-dependence and reentrance behavior observed in simulations.

## 4 Two populations

In Sec. 3.1, we have demonstrated that a neural population with strong excitatory recurrent connections supports bistability. For a network of *N* weakly-coupled bistable units, there are 3^*N*^ fixed points with 2^*N*^ stable nodes and many saddle points. These fixed points define a “landscape” for the dynamics. In the limit *N* → ∞, the competition between self-coupling and cross-coupling even gives rise to transient chaos [38]. In this section, we briefly discuss the general case of *N* populations and then focus on two populations. We will see how synaptic depression, as well as cross-coupling, facilitate sequences of state transitions in response to uniform stimuli.

### 4.1 Fixed points and stability

In general, for a network of *N* bistable units [Eqs. (4)–(6)], fixed points are solutions to a set of nonlinear equations (*i* = 1, …, *N*),

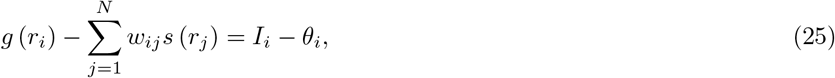

where 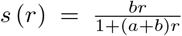 is the steady synaptic current. Once we find a solution *r*\* = (*r*_1_, …, *r_N_*)^*T*^, we can get the values of *s* and *d* at the steady-state from Eq. (11). Linearizing the flow near a fixed point yields a 3*N* × 3*N* Jacobian matrix with 3N eigenvalues {Λ_*i*_}. For a hyperbolic fixed point (Reλ_*i*_ ≠ 0, ∀*i*), the local stability is captured by these eigenvalues: The fixed point is a stable node if Reλ_*i*_ < 0 for all *i*, and it is a saddle if Reλ_*j*_ > 0 for some (but not all) *j*. If there are *k* eigenvalues with positive real parts, we refer to this fixed point as a *k*-saddle (A stable node is a 0-saddle).

Heuristically, for *N* weakly interacting bistable units, each has two stable nodes and one 1-saddle. The number of *k*-saddles is

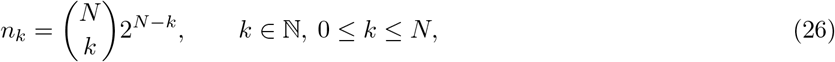

which amounts to choosing *k* positive real eigenvalues out of *N* eigenvalues then multiplying the number of remaining (*N* – *k*) bistable states. It is straightforward to check that 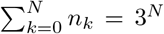. The number of saddles grows exponentially as the network size increases, as does the number of stable nodes. We should note that this estimation works only in the limit of weak cross-coupling. However, there can be still a large number of stable fixed points and even more saddles in a coupled network.

### 4.2 Linearization for *N* = 2

For two populations, if we denote a fixed point as (*r*_1_, *s*_1_, *d*_1_, *r*_2_, *s*_2_, *d*_2_)^*T*^, the Jacobian can be written as blocked matrices,

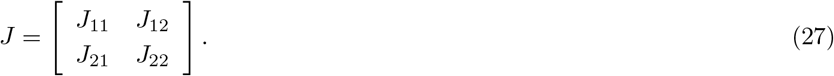

The 3 × 3 matrix *J_ij_* (*i*, *j* = 1, 2) is evaluated at the fixed point (*δ_ij_* is the Kronecker delta)

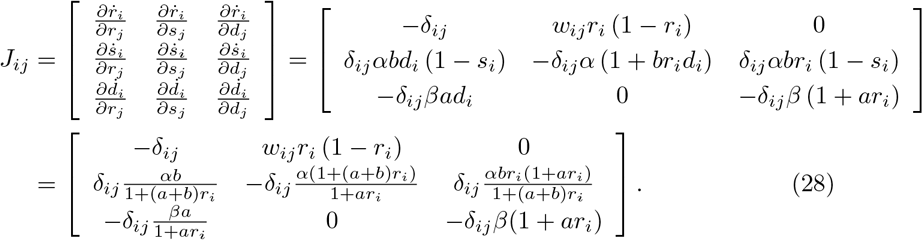

#### Basins of attraction

It is straightforward to compute all six eigenvalues of *J* and examine how many of them have negative real parts. For instance, if there is no cross-coupling (*w_ij_* = 0), there should be five saddles and four stable nodes. The stable nodes correspond to: Both units are OFF (“00”); Both are ON (“11”); One is OFF, and the other is ON (“01” and “10”). For any given initial condition, the system approaches one of the four possible stable nodes as its final stable state. The likelihood of landing on a specific node depends on the size of its basin of attraction, which may differ quite a lot between nodes.

Weak coupling (*w_ij_* ≪ *w_ii_*) does not change the number of fixed points but can alter their stability as well as the number and sizes of basins of attraction. In Fig. 6, we show the fixed points of two symmetrically coupled units (*w*_12_ = *w*_21_) without external stimuli and their basins of attraction projected in the *r*_1_-*r*_2_ plane. ^[1]^ Excitatory coupling [Fig. 6(b)] enlarges the basin of the (11) state while mutual inhibition [Fig. 6(c), (d)] destabilizes the state. Its area shrinks quickly upon receiving a small amount of inhibition. When *w*_12_ = *w*_21_ = −0.5, the (11) state becomes a saddle. Basins of attraction for the other three stable states deform into complex structures near the (11) state. So the final state depends sensitively on the properties of the stimulus and, we shall see later, upon the initial condition.

**Figure 6.**
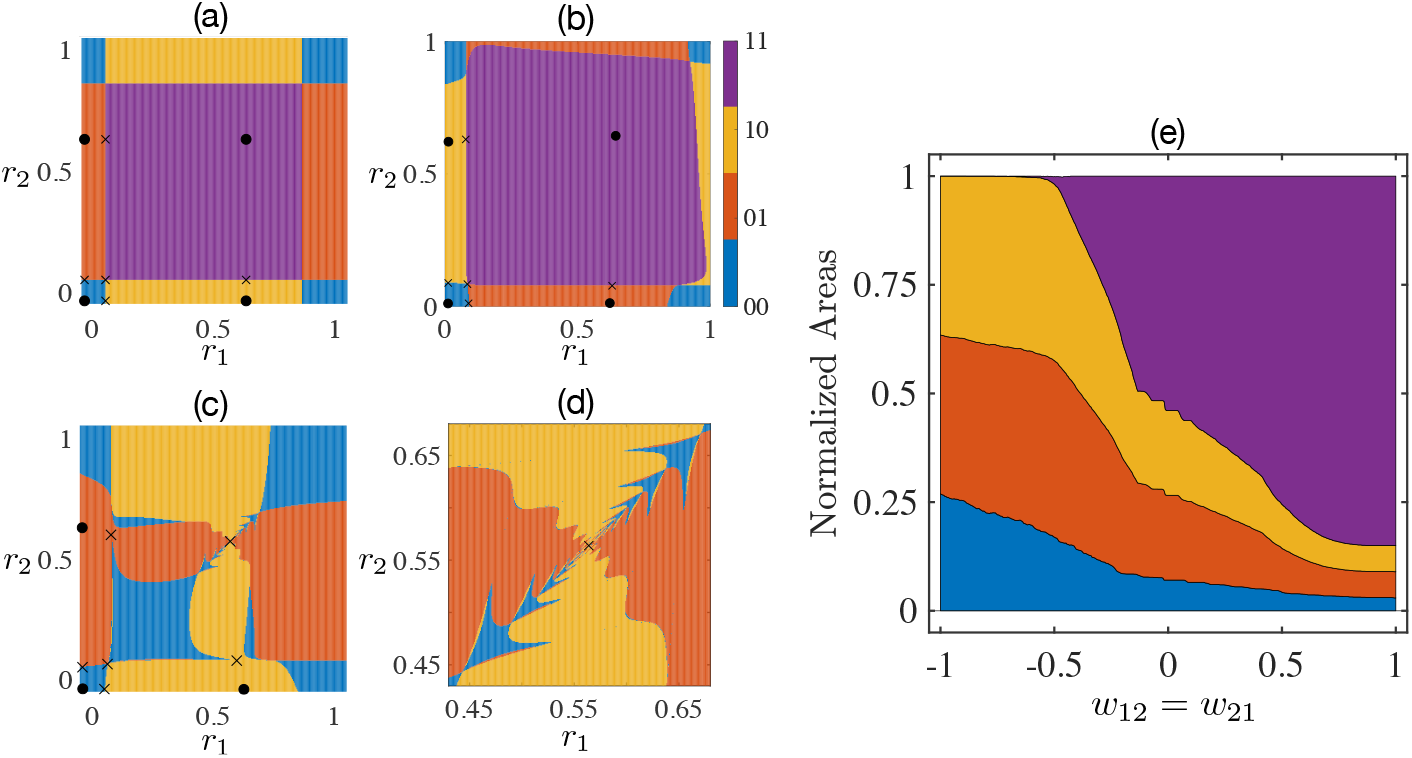
Fixed points and projected basins of attraction for two symmetrically coupled units (*w*_12_ = *w*_21_). (a) – (d): Stable nodes (saddles) are labeled as filled circles (crosses) and surround by color-coded basins. The self-coupling is fixed at *w*_11_ = *w*_22_ = 40 and cross-couplings are chosen as a)*w*_12_ = 0, b) *w*_12_ = 0.5, c)-d) *w*_12_ = −1. (d) Zoom-in details reveal fine structures near the (11) state. (e) Normalized basin areas of stable attractors as a function of the cross-coupling.

#### Number of Fixed Points and Bifurcations

To further examine the effects of cross-coupling between two identical populations, we apply numerical continuation on the stable states by varying *w_ij_* and monitoring the eigenvalues of the Jacobian. Although we are mainly interested in weak coupling, we extend the search to fairly large |*w_ij_* | for completeness. For moderate coupling strengths, the system can support up to four stable fixed points. Too much excitation eliminates low-rate states, leaving both units on. Figure 7 shows regions associated with distinct sets of stable fixed points in the *w*_12_-*w*_21_ plane. Inset graphs (a – f) are schematic illustrations of the configuration of fixed points under different pairs of cross-coupling strength (*w*_12_, *w*_21_). Traversing between different regions in the main panel (illustrated by the correspondingly labeled inset graphs) leads to saddle-node (*b* → *c* and *a* → *d*) or Hopf bifurcations (*a* → *b* and *d* → *e*). Within the same region, there are bifurcations that change the stability of saddles (*d* → *f*). Because of the existence of higher-order saddles, it is worth noting that the figure is merely a glimpse of the complex dynamics in a six-dimensional space.

**Figure 7.**
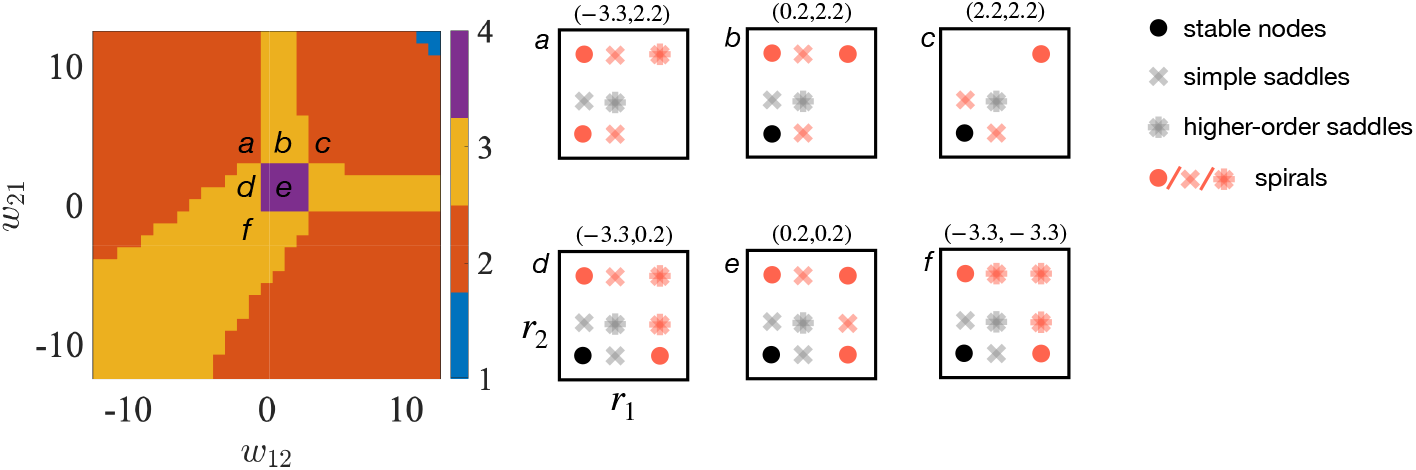
Effects of cross-coupling on the number of stable fixed points for two identical populations. The labels (a-f) indicate the parameters of the corresponding inset panels (a – f), which are schematics of the system’s fixed points with different pairs of cross-coupling strength (*w*_12_, *w*_21_). In the inset panels, solid dots are stable fixed points; crosses are simple saddles; stars denote higher-order saddles with more than one positive eigenvalue. Red symbols indicate the existence of complex eigenvalues.

### 4.3 Responses to stimuli

Similarly to the single population case, two weakly-coupled populations generate rich responses to stimuli, especially if the populations are not identical and are coupled asymmetrically. Figure 8 shows numerical results of various state transitions in a two-population system with weak cross-inhibition. Each population is initially OFF [in the (00) state] and has different input thresholds. A stimulus, which provides equal input to the two populations, may temporarily activate one or both populations. When the stimulus is over, the system approaches a steady state. As in previous sections, the final state reached by the system depends on the amplitude and duration of received stimuli. In panel (a) of Fig. 8, increasing duration leads to a sequence of four possible final states as the trajectory of network activity traverses the basins of attraction (00) → (10) → (11) → (01), with the final state corresponding to the basin upon stimulus offset. In panel (b), with increasing amplitude, the corresponding state-transitions are: (00) → (10) → (01) → (11) → (10).

**Figure 8.**
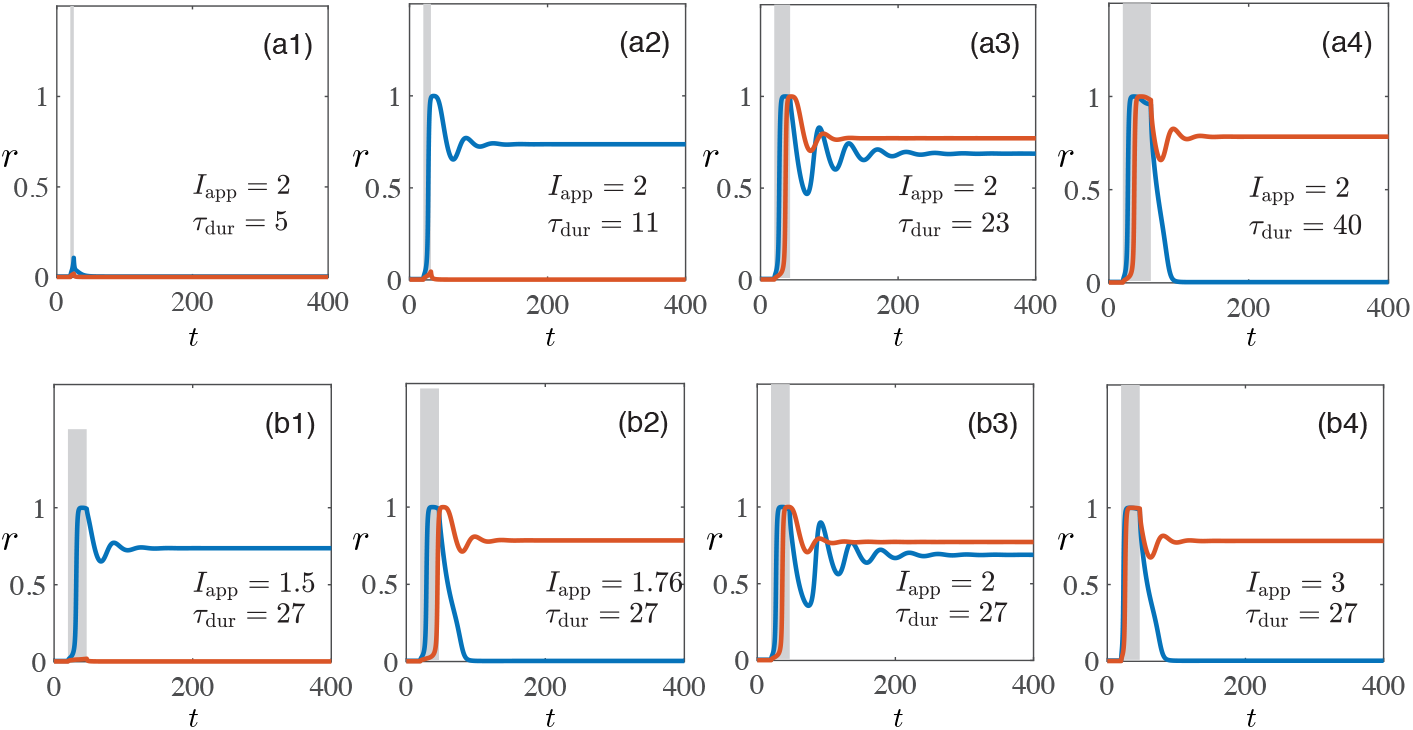
Responses of two populations to stimuli (gray bars). Blue and red lines denote population 1 and 2 respectively. (a1)-(a4) Stimuli with a fixed amplitude (*I*_app_ = 2) and increasing duration. (b1)-(b4) Stimuli with a fixed duration (*τ*_dur_ = 27) and increasing amplitude. Both panels show re-entrance behavior. The coupling weights are *w*_11_ = 47, *w*_12_ = −1.2, *w*_21_ = −0.4, *w*_22_ = 54. The thresholds are *θ*_1_ = 5.6 and *θ*_2_ = 6.4.

If we choose different initial states to start with, the transitions and the final states are even richer. Figure 9 summarizes steady-states for four initial conditions [(00), (01), (10), (11)] as functions of the stimuli duration and amplitude. Some of the phase boundaries resemble those seen in a single population [Fig. 4], but the type and number of final states are different. For example, if the initial state is (10), then after uniform stimulation, the system either remains in this state or switches to the (01) state or the (11) state with strong and short input. If we start with the (01) state, then the system may go to the (11) state or switch to the (10) state, but never return to the initial state with strong input. Such history-dependent responses can be used to encode stimulus propeties in memory.

**Figure 9.**
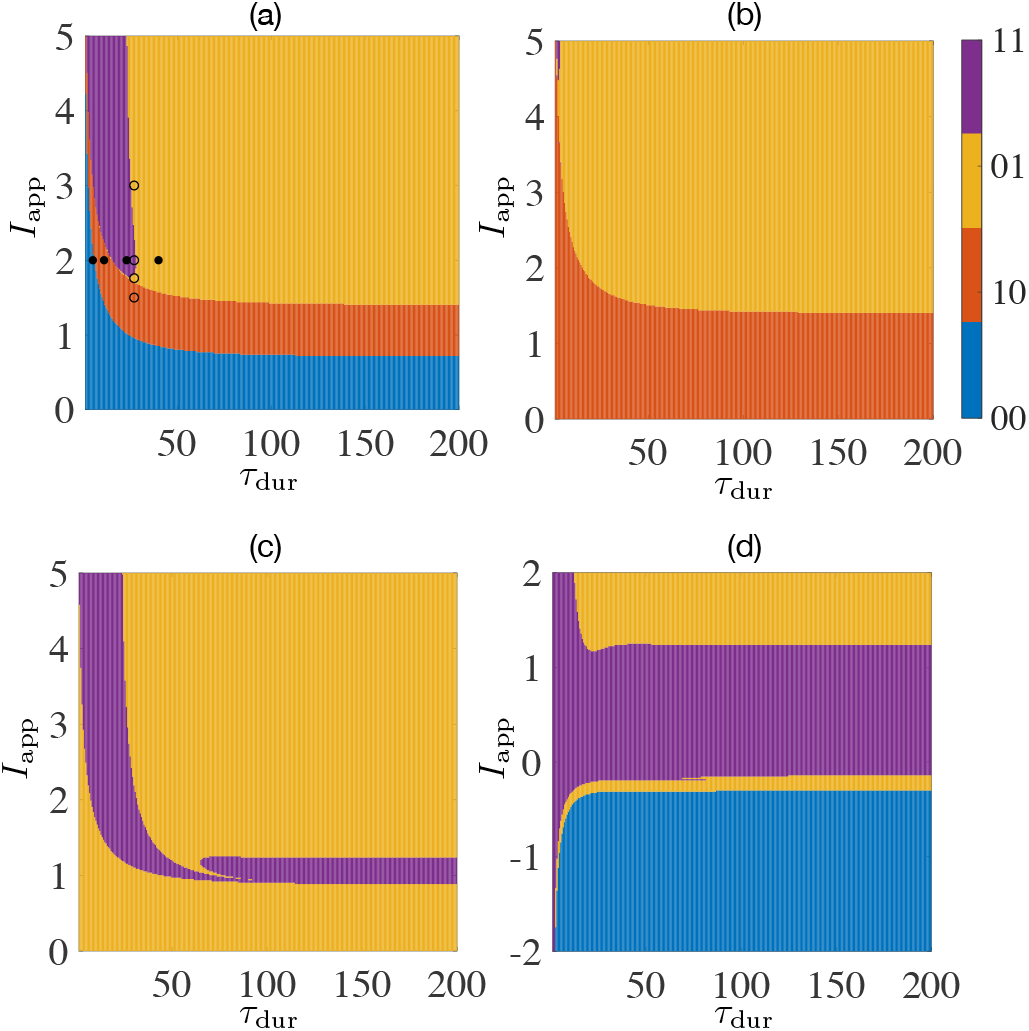
Phase diagrams for two populations with asymmetric cross-inhibition. (a) – (d): Initial states are (a) (00); (b) (10); (c) (01); and (d) (11). The filled and open circles in panel (a) correspond to subplots (a) and (b) in Fig. 8.

## 5 Multiple populations

As we have seen in the two population case, weak cross-coupling has a noticeable influence on the system’s dynamics. Previous sections also demonstrate that recurrent inputs may change the stability of fixed points. In a typical network with weak cross coupling, the number of stable fixed points is usually less than 2^*N*^. The remaining attractors and saddles allow rich transient dynamics under stimuli that lead to itinerancy in the state space.

We numerically explore the responses of multiple neural populations to transient stimuli that are uniform across the network (that is, the same input to all populations). Parameters including the self-coupling strengths are chosen such that each population is bistable. Cross coupling strengths in the weight matrix are drawn from a Gaussian distribution 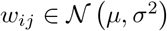, *i* ≠ *j*. Before a stimulus, the system is at a steady state (an attractor). After the stimulus, the system either remains in the initial state or settles in another state. For different stimuli applied to a specific network, the number of final states that are reachable with a uniform stimulus measures the capacity of the network to encode features such as the amplitude and duration of the stimulus via attractor state itinerancy (or state transitions).

For small networks, we can identify all fixed points (nodes and saddles) and track the state transitions. Through three examples, Figs. 10–12 we illustrate how connectivity and initial conditions affect a network’s dynamical response to uniform stimuli. In each case, we run multiple trials with random weight matrices. For each trial, we find all the stable fixed points and start the simulation from a specific one. Applying the same stimulation protocol to all units (square-wave currents with varying amplitudes and durations), we count the number of distinct steady states after the simulation (i.e., the number of reachable final states). Then we plot trial-averaged results against the mean *μ* and the standard deviation *σ* of non-diagonal entries in the weight matrix.

**Figure 10.**
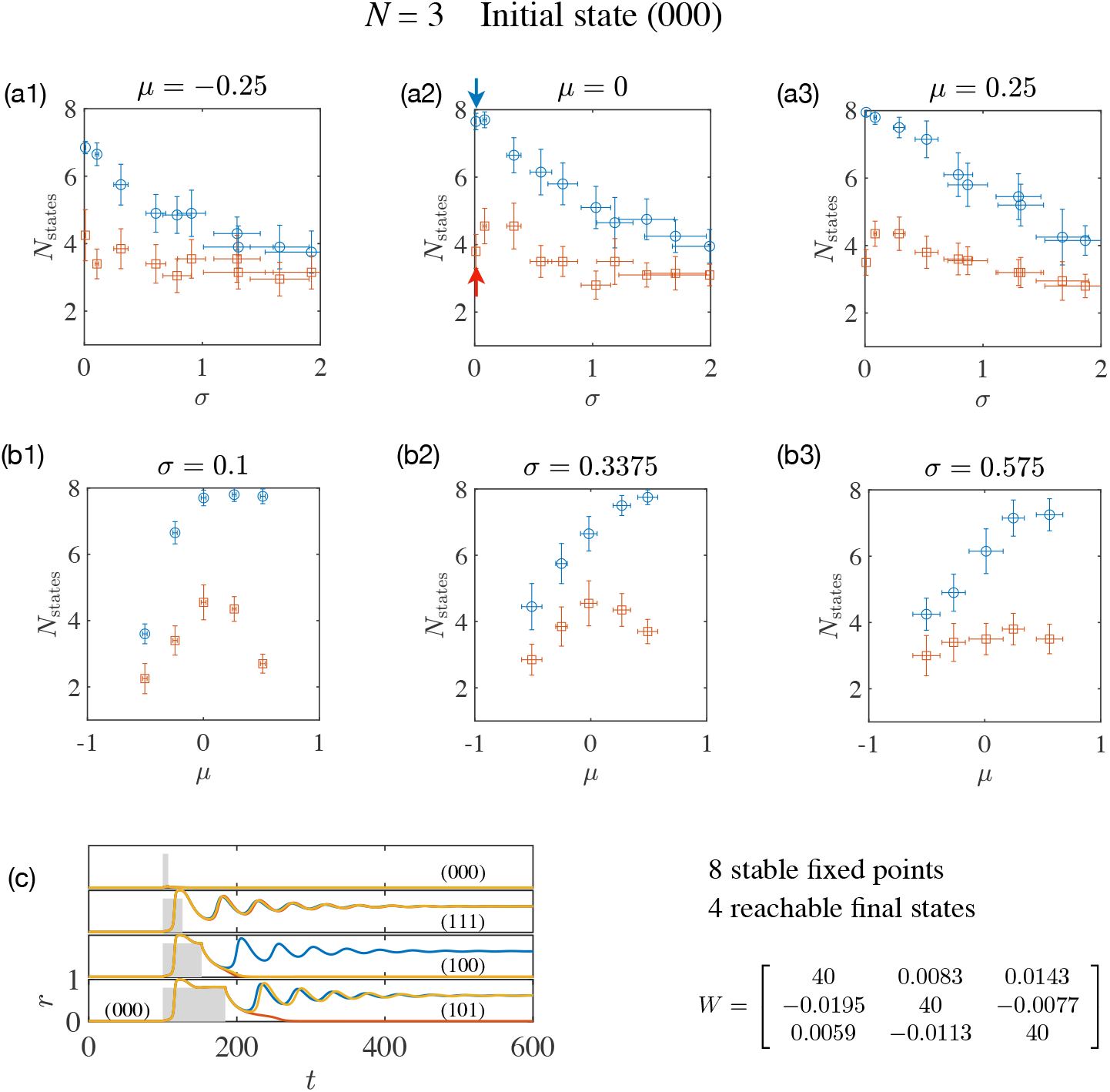
Numbers of attractors (blue open circles) and reachable final states (red open squares) for a three-population network with initial state (000). In panels (a) and (b), results are plotted with varying mean values *μ* and standard deviations *σ* of cross couplings in the weight matrix *W*. Panel (c) shows firing rates before and after stimuli (gray rectangular bars) for the network indicated by blue and red arrows in (a2). Digits in parentheses indicate the distributed activity where 1(0) denotes the ON (OFF) state for each unit.

**Figure 11.**
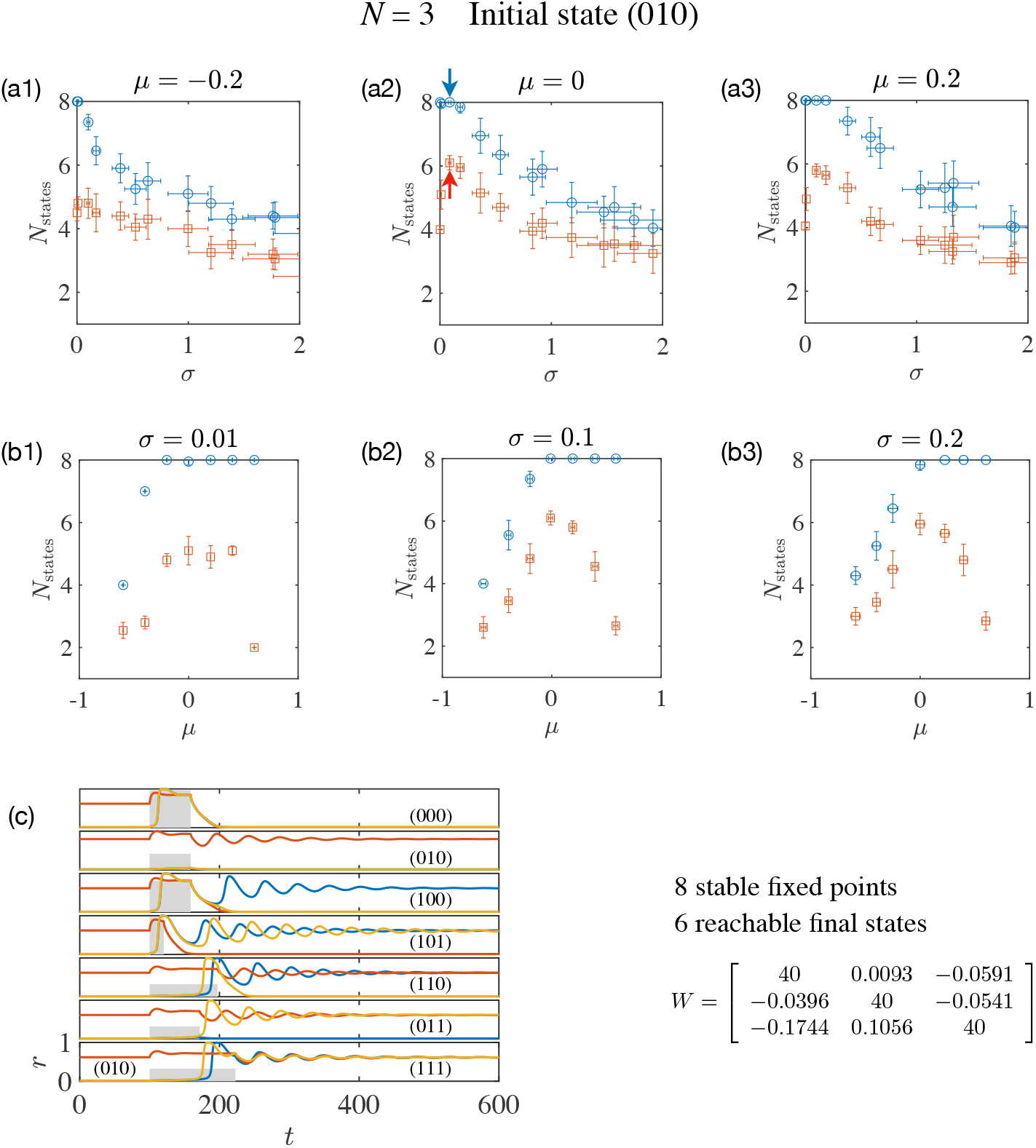
Numbers of attractors (blue open circles) and reachable final states (red open squares) for a three-population network with initial state (010). Labels are similar to those defined in Fig. 10.

**Figure 12.**
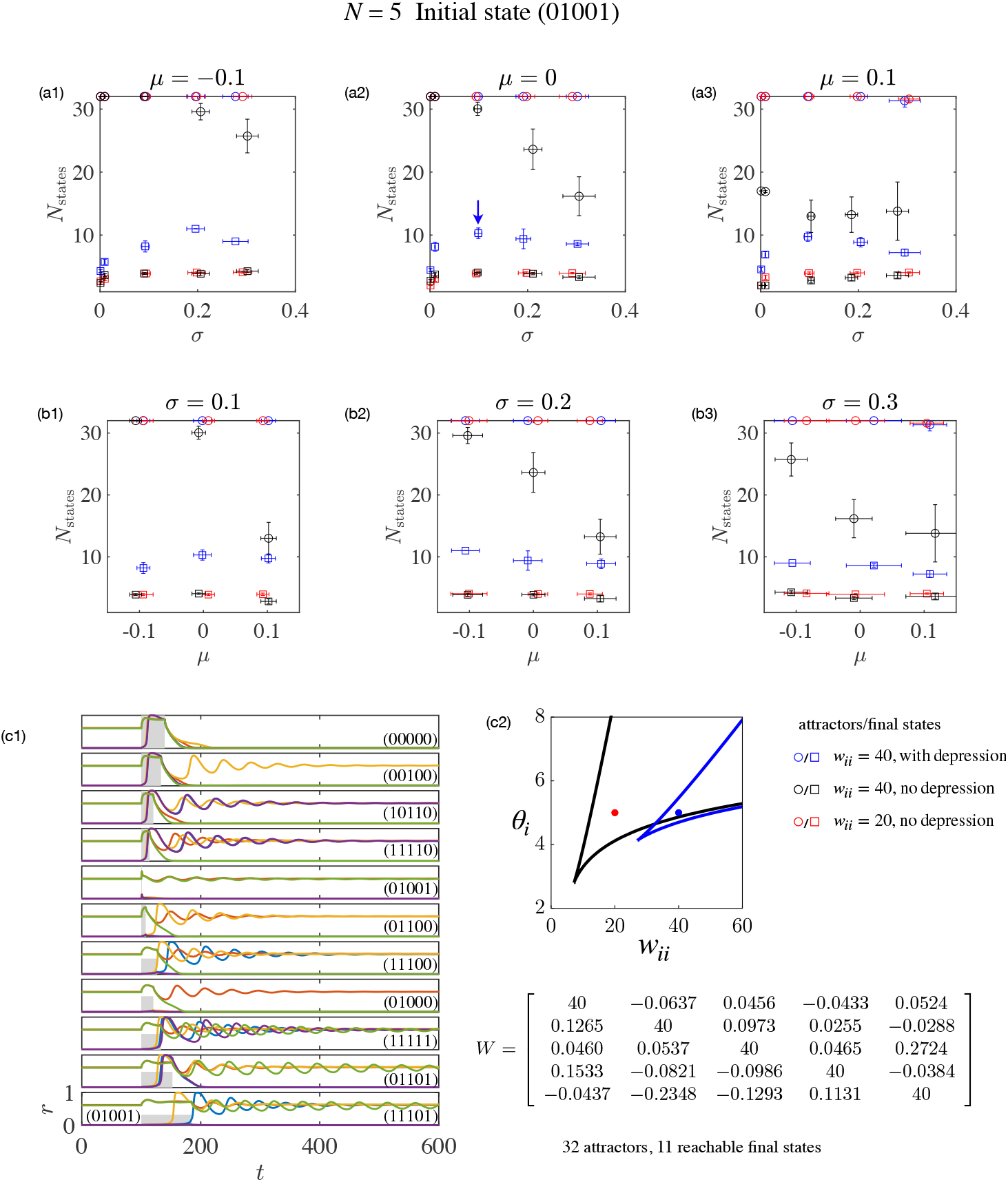
Numbers of attractors (blue open circles) and reachable final states (red open circles) for a five-population network with initial state (01001). Labels are similar to those defined in Fig. 10, except for the color-coding, which indicates three cases with and without depression. For a specific weight matrix *W* (indicated by a blue arrow in (a2)), panel (c1) shows the time evolution as well as 11 distinct final states. Panel (c2) shows the bistable regions with (blue lines) and without (black lines) synaptic depression. The blue point locates at (*w_ii_* = 40, *θ_i_* = 5) and red points stands for (*w_ii_* = 20, *θ_i_* = 5).

In Fig. 10, we set *N* = 3 and choose (000) as the initial state. With fixed mean cross-coupling strength, *μ*, panels (a1) – (a3) show the number of attractors (open circles) and the number of final states reachable following a transient, uniform stimulation (open squares) as functions of the standard deviation of the cross-coupling strengths, *σ*. Although the network naturally has fewer attractors as *σ* increases, the number of reachable final states show non-monotonic behavior. When excitation balances or exceeds inhibition (a2 and a3), weak random couplings facilitate state transitions. The number of reachable states increases and maximizes at an optimal value of *σ*. When inhibition surpasses excitation (a1), variance in *w_ij_* smooths out the peak.

In panels (b1) – (b3), we fix the standard deviation of the cross-coupling strength, *σ*, and show how the numbers of states scale with changing the mean cross-coupling strength, *μ*. When the randomness in the couplings is low (b1), the number of attractors approaches 2^*N*^ for *μ* ≥ 0. The number of reachable states peaks around *μ* = 0 (as in b1 and b2). So the network’s optimal performance requires excitation-inhibition balance and weak (but non-zero) random couplings. For large randomness (b3), there are fewer stable states so that the number of reachable states is small (≈3).

In panel (c), the stacked plots show firing rates in response to uniform transient stimulations. Starting with initial state (000) and very weak heterogeneity in the random cross-coupling [*σ* = 0.01, *μ* = 0 as indicated by arrow in (a2)], the network has eight attractor states. Following uniform stimulations which differ in their amplitudes and durations, there are four distinct final states: (000), (111), (100) and (101).

Figure 11 demonstrates the effect of initial conditions on the dynamics of a three-population network. Here we choose (010) as the initial state. Compared with Fig. 10, in response to uniform stimuli the network with weak random couplings visits more distinct states (a2 and a3). Panel (c) shows the optimal case (*σ* = 0.1) where the system supports all eight attractors and reaches seven of them, that is all except for (001).

In Fig. 12, we compare the performance of a five-unit network with and without synaptic depression. Panels (a) and (b) show the number of attractors (open circles) and the number of reachable final states (open squares). The color-coding corresponds to three cases:

i. *w_ii_* = 40 with depression (blue) Similar to the three-unit case (Figs. 10 & 11), the network supports 2^5^ = 32 attractors when the random cross-coupling is weak. The number of reachable final states peaks when both the mean cross-coupling strength, *μ*, is small or zero (b1-b3) and the standard deviation, *σ* is small but non-zero (a1-a3). For instance, as indicated in by a blue arrow in subplot (a,2), there are 11 reachable final states at *μ* = 0 and *σ* = 0.1. The detailed time evolutions for an example weight matrix are shown in panel (c1).
ii. *w_ii_* = 40 without depression (black) When we fix the depression variable at unity, *d* = 1, and keep all other parameters unchanged, the network reaches much fewer final states (≈ 3). As shown in subplot (a3), a small mean strength of cross excitation (*μ* = 0.1) greatly reduces the total number of attractors. This can be understood by looking at panel (c2) where we graph the bistable region in the *w_ii_*-*θ_i_* plane. The blue dot indicates the chosen parameters as (*w_ii_* = 40, *θ_i_* =5). It is within the bistable region (blue lines) but is located near the lower boundary (black lines). Therefore, a small excitatory input can drive a unit out of the bistable region. When some units are no longer bistable, the total number of attractors become much less than 32.
iii. *w_ii_* = 20 without depression (red) In the absence of depression, if we choose parameters in the middle of the bistable region as (*w_ii_* = 20, *θ_i_* = 5) indicated by a red dot, the network has 32 attractors but much fewer reachable final states.

As these two control groups show, synaptic depression indeed increases the number of state-transitions.

## 6 Discussion

In this paper, we consider small circuits of bistable neural populations with synaptic depression, focusing on the circuit’s responses to uniform stimuli with different amplitudes and durations. Because of the multiple time-scales between firing rates, synaptic currents, and depression, the system is near several bifurcations that may deform the basins of attraction dramatically. The final state of the system after a perturbation thus sensitively depends on the properties of the perturbing stimulus.

In the absence of cross-coupling, the number of stable fixed points of the system is 2^*N*^, where *N* is the number of bistable units. While the number of stable fixed points is maximized in this limit, the lack of interaction between units means the responses to stimuli and the history dependence is rather limited. Conversely, with very strong cross-couplings, subsets of units become very highly correlated in their activity, reducing the effective *N*: for example, two units with strong reciprocal cross-excitation are always ON together or OFF together, so act together more like a single unit. We find that with weak cross-couplings, the total number of stable fixed points can remain high, while the interactions between units enables a simple, uniform stimulus (identical to all units) to cause a network response that traces a high-dimensional trajectory through the space of units’ activities. The high-dimensionality of the response leads to history-dependence and a richness in the types of stable states achievable by a stimulus that excites all units equally. This behavior allows networks of many units to retain separate information about the amplitude, duration, and number of identical, repeated stimuli[24, 32].

Our work follows that of others demonstrating the richness of states in networks with coupled units. Prior work showed that in the macroscopic limit, with weak self-coupling and strong, balanced cross-coupling, a chaotic regime exists [39], whereas when the self-coupling is strong enough that each unit is bistable, multiple stable states exist and can be reached by transient chaos [38]. Here, we focused on smaller circuits and included the impact of synaptic depression, a common feature of cortical synapses. Synaptic depression can reduce the total number of fixed points by reducing the stability of the ON state (active synapses are effectively weakened by depression). However, the same effect can enhance the number of states reachable by a uniform stimulus, as a weakening of the connections within previously active units allows new units to become ON when the duration of the stimulus is extended. Similarly, such relative destabilization of previously active states enhances the history dependence of stimulus responses and causes the network’s activity to explore a wider range of the state space. We expect that incorporation of firing-rate adaptation in the neural responses would have a similar effect in destabilizing active states.

The dependence of network activity on the duration of stimuli or interval between stimuli is particularly noticeable when intervals on the order of a few hundred milliseconds are present in auditory tasks. Synaptic depression operates on a suitable time scale to produce the ongoing network dynamics that could account for such interval or duration dependence [40].

While our work here focuses on the dynamics of network behavior in the presence of a stimulus which is constant in time, the dependence on initial conditions of the network’s response to a given stimulus (Fig. 10 and Fig. 11) imbues the network with history-dependence. Therefore, the network can respond differently, according to the number and/or types of and/or order of preceding stimuli[24, 32, 40]. In this manner, such networks could account for the observed transitions of neural activity through a set of distinct attractor states during a counting task [32, 41] and could even provide a basis for context-dependent integration of stimulus properties[42].

## List of abbreviations

HB: Hopf Bifurcation
LP: Limit Point
SN: Saddle-Node
SHO: Saddle-Homoclinic Orbit

## Ethics approval and consent to participate

Not applicable.

## Consent for publication

Not applicable.

## Availability of data and materials

The datasets generated and/or analysed during the current study are available at https://github.com/blchen00/attractor-itinerancy-paper/.

## Author’s contributions

PM designed the project. BC carried out the analysis. Both authors wrote this paper, read and approved the final manuscript.

## Acknowledgements

BC acknowledges the financial support from the Swartz Foundation, as well as helpful conversations with Benjamin Ballintyn.

## Footnotes

[1] We choose a rate vector *r*_0_ = (*r*_1_, *r*_2_)^*T*^ and set the initial condition as (*r*_0_, *s*(*r*_0_), *d*(*r*_0_)), then label the final state after integrating the differential equations for a long time.

## References

1. Snowdon, C.T.: Response of nonhuman animals to speech and to species-specific sounds. Brain, behavior and evolution 16(5-6), 409–429 (1979)

2. Fuster, J.M., Jervey, J.P.: Inferotemporal neurons distinguish and retain behaviorally relevant features of visual stimuli. Science 212(4497), 952–955 (1981)

3. Funahashi, S., Bruce, C.J., Goldman-Rakic, P.S.: Mnemonic coding of visual space in the monkey’s dorsolateral prefrontal cortex. Journal of neurophysiology 61(2), 331–349 (1989)

4. Sigala, N., Logothetis, N.K.: Visual categorization shapes feature selectivity in the primate temporal cortex. Nature 415(6869), 318 (2002)

5. Leutgeb, J.K., Leutgeb, S., Treves, A., Meyer, R., Barnes, C.A., McNaughton, B.L., Moser, M.-B., Moser, E.I.: Progressive transformation of hippocampal neuronal representations in “morphed” environments. Neuron 48(2), 345–358 (2005)

6. Rotshtein, P., Henson, R.N., Treves, A., Driver, J., Dolan, R.J.: Morphing marilyn into maggie dissociates physical and identity face representations in the brain. Nature neuroscience 8(1), 107 (2005)

7. Daelli, V., Treves, A.: Neural attractor dynamics in object recognition. Experimental brain research 203(2), 241–248 (2010)

8. Miller, P.: Itinerancy between attractor states in neural systems. Current opinion in neurobiology 40, 14–22 (2016)

9. Deppisch, J., Pawelzik, K., Geisel, T.: Uncovering the synchronization dynamics from correlated neuronal activity quantifies assembly formation. Biological cybernetics 71(5), 387–399 (1994)

10. Radons, G., Becker, J., Dülfer, B., Kruger, J.: Analysis, classification, and coding of multielectrode spike trains with hidden markov models. Biological cybernetics 71(4), 359–373 (1994)

11. Gat, I., Tishby, N., Abeles, M.: Hidden markov modelling of simultaneously recorded cells in the associative cortex of behaving monkeys. Network: Computation in neural systems 8(3), 297–322 (1997)

12. Otterpohl, J., Haynes, J., Emmert-Streib, F., Vetter, G., Pawelzik, K.: Extracting the dynamics of perceptual switching from ‘noisy’ behaviour: An application of hidden markov modelling to pecking data from pigeons. Journal of Physiology-Paris 94(5-6), 555–567 (2000)

13. Rainer, G., Miller, E.K.: Neural ensemble states in prefrontal cortex identified using a hidden markov model with a modified em algorithm. Neurocomputing 32, 961–966 (2000)

14. Jones, L.M., Fontanini, A., Sadacca, B.F., Miller, P., Katz, D.B.: Natural stimuli evoke dynamic sequences of states in sensory cortical ensembles. Proceedings of the National Academy of Sciences 104(47), 18772–18777 (2007)

15. Escola, S., Fontanini, A., Katz, D., Paninski, L.: Hidden markov models for the stimulus-response relationships of multistate neural systems. Neural computation 23(5), 1071–1132 (2011)

16. Abeles, M., Bergman, H., Gat, I., Meilijson, I., Seidemann, E., Tishby, N., Vaadia, E.: Cortical activity flips among quasi-stationary states. Proceedings of the National Academy of Sciences 92(19), 8616–8620 (1995)

17. Latimer, K.W., Yates, J.L., Meister, M.L., Huk, A.C., Pillow, J.W.: Single-trial spike trains in parietal cortex reveal discrete steps during decision-making. Science 349(6244), 184–187 (2015)

18. Miller, P., Katz, D.B.: Stochastic transitions between neural states in taste processing and decision-making. Journal of Neuroscience 30(7), 2559–2570 (2010)

19. Litwin-Kumar, A., Doiron, B.: Slow dynamics and high variability in balanced cortical networks with clustered connections. Nature neuroscience 15(11), 1498 (2012)

20. Miller, P., Katz, D.B.: Accuracy and response-time distributions for decision-making: linear perfect integrators versus nonlinear attractor-based neural circuits. Journal of computational neuroscience 35(3), 261–294 (2013)

21. Doiron, B., Litwin-Kumar, A.: Balanced neural architecture and the idling brain. Frontiers in computational neuroscience 8, 56 (2014)

22. Ashwin, P., Creaser, J., Tsaneva-Atanasova, K.: Sequential escapes: onset of slow domino regime via a saddle connection. The European Physical Journal Special Topics 227(10-11), 1091–1100 (2018)

23. Kilpatrick, Z.P., Bressloff, P.C.: Binocular rivalry in a competitive neural network with synaptic depression. SIAM Journal on Applied Dynamical Systems 9(4), 1303–1347 (2010)

24. Miller, P.: Stimulus number, duration and intensity encoding in randomly connected attractor networks with synaptic depression. Frontiers in computational neuroscience 7, 59 (2013)

25. Moreno-Bote, R., Rinzel, J., Rubin, N.: Noise-induced alternations in an attractor network model of perceptual bistability. Journal of Neurophysiology 98(3), 1125–1139 (2007)

26. Shpiro, A., Moreno-Bote, R., Rubin, N., Rinzel, J.: Balance between noise and adaptation in competition models of perceptual bistability. Journal of computational neuroscience 27(1), 37 (2009)

27. Tsodyks, M.V., Markram, H.: The neural code between neocortical pyramidal neurons depends on neurotransmitter release probability. Proceedings of the National Academy of Sciences 94(2), 719–723 (1997)

28. Varela, J.A., Sen, K., Gibson, J., Fost, J., Abbott, L., Nelson, S.B.: A quantitative description of short-term plasticity at excitatory synapses in layer 2/3 of rat primary visual cortex. Journal of Neuroscience 17(20), 7926–7940 (1997)

29. Tsodyks, M., Pawelzik, K., Markram, H.: Neural networks with dynamic synapses. Neural computation 10(4), 821–835 (1998)

30. Tabak, J., Senn, W., O’Donovan, M.J., Rinzel, J.: Modeling of spontaneous activity in developing spinal cord using activity-dependent depression in an excitatory network. Journal of Neuroscience 20(8), 3041–3056 (2000)

31. Wilson, H.R., Cowan, J.D.: Excitatory and inhibitory interactions in localized populations of model neurons. Biophysical journal 12(1), 1–24 (1972)

32. Ballintyn, B., Shlaer, B., Miller, P.: Discrimination and recall of sequences of stimuli in randomly connected attractor networks with short-term synaptic depression. Journal of Computational Neuroscience, (2019)

33. Beer, R.D.: On the dynamics of small continuous-time recurrent neural networks. Adaptive Behavior 3(4), 469–509 (1995)

34. Ermentrout, G.B., Terman, D.H.: Mathematical Foundations of Neuroscience vol. 35, 1st edn. Springer, New York (2010)

35. Nan, P., Wang, Y., Kirk, V., Rubin, J.E.: Understanding and distinguishing three-time-scale oscillations: Case study in a coupled morris–lecar system. SIAM Journal on Applied Dynamical Systems 14(3), 1518–1557 (2015)

36. Rubin, J., Terman, D.: Analysis of clustered firing patterns in synaptically coupled networks of oscillators. Journal of mathematical biology 41(6), 513–545 (2000)

37. Terman, D., Ahn, S., Wang, X., Just, W.: Reducing neuronal networks to discrete dynamics. Physica D: Nonlinear Phenomena 237(3), 324–338 (2008)

38. Stern, M., Sompolinsky, H., Abbott, L.: Dynamics of random neural networks with bistable units. Physical Review E 90(6), 062710 (2014)

39. Van Vreeswijk, C., Sompolinsky, H.: Chaos in neuronal networks with balanced excitatory and inhibitory activity. Science 274(5293), 1724–1726 (1996)

40. Goudar, V., Buonomano, D.V.: A model of order-selectivity based on dynamic changes in the balance of excitation and inhibition produced by short-term synaptic plasticity. American Journal of Physiology-Heart and Circulatory Physiology (2014)

41. Morcos, A.S., Harvey, C.D.: History-dependent variability in population dynamics during evidence accumulation in cortex. Nature neuroscience 19(12), 1672 (2016)

42. Mante, V., Sussillo, D., Shenoy, K.V., Newsome, W.T.: Context-dependent computation by recurrent dynamics in prefrontal cortex. nature 503(7474), 78 (2013)

